# Biomechanics of the peafowl’s crest reveals frequencies tuned to social displays

**DOI:** 10.1101/197525

**Authors:** Suzanne Amador Kane, Daniel Van Beveren, Roslyn Dakin

## Abstract

Feathers act as vibrotactile sensors that can detect mechanical stimuli during avian flight and tactile navigation, suggesting that they may also detect stimuli during social displays. In this study, we present the first measurements of the biomechanical properties of the feather crests found on the heads of birds, with an emphasis on those from the Indian peafowl (*Pavo cristatus*). We show that in peafowl these crest feathers are coupled to filoplumes, small feathers known to function as mechanosensors. We also determined that airborne stimuli with the frequencies used during peafowl courtship and social displays couple efficiently via resonance to the vibrational response of their feather crests. Specifically, vibrational measurements showed that although different types of feathers have a wide range of fundamental resonant frequencies, peafowl crests are driven near-optimally by the shaking frequencies used by peacocks performing train-rattling displays. Peafowl crests were also driven to vibrate near resonance in a playback experiment that mimicked the effect of these mechanical sounds in the acoustic very near-field, reproducing the way peafowl displays are experienced at distances ≤ 1.5m *in vivo*. When peacock wing-shaking courtship behaviour was simulated in the laboratory, the resulting airflow excited measurable vibrations of crest feathers. These results demonstrate that peafowl crests have mechanical properties that allow them to respond to airborne stimuli at the frequencies typical of this species’ social displays. This suggests a new hypothesis that mechanosensory stimuli could complement acoustic and visual perception and/or proprioception of social displays in peafowl and other bird species. We suggest behavioral studies to explore these ideas and their functional implications.

## Introduction

Bird feathers are known to act as mechanosensors that allow birds to detect and respond to a variety of mechanical stimuli [1–3]. For example, flight, contour, and facial bristle feathers can act as sensors that provide important information during flight and prey capture [4,5,3,6,7]. Indeed, feathers have been suggested to have evolved originally to serve sensory functions, because even isolated protofeathers could have played a sensory role before the evolution of specialized arrays of feathers that enabled flight or thermoregulation [8]. Feather head crests have been found in fossils of some dinosaurs and early birds as well as a wide variety of living species of birds [9,10]. While feather crests have usually been studied for their roles as possible visual signals [11–13, see also 14 for a review], recent behavioral studies of two auklet species have shown that their erect head crest feathers can play a mechanosensory role during tactile navigation similar to that of mammalian whiskers and arthropod antennae [14,15]. These findings suggest that feather crests in other birds also might play a previously unrecognized mechanosensory functional role. Crest feathers and other types of feathers found on the heads of birds have received little attention in the literature, especially in comparison to the significant body of literature on the morphology and mechanical properties of wing, tail and train covert feathers [16]. In addition, no research has considered whether birds, like some arthropods, might detect air-borne stimuli generated during social displays via mechanoreception, or what influence this may have on their social interactions.

Here we report on the first biomechanical study of bird crest feathers, with an emphasis on understanding how their physical properties might relate to their various possible functions. This work focused on the large crest of the Indian Peafowl (*Pavo cristatus*), which is found on both sexes [17]. The male of this species (“peacock”) performs elaborate multimodal courtship displays accompanied by mechanical sound experienced by nearby females. During their displays, male Indian peafowl (“peacocks”) attract mates by spreading and erecting their train (a fan-like array of long, colorful feathers) and performing two dynamic courtship behaviors. First, during “wing-shaking, the male flaps his partially-unfurled wings at approximately 5.4 Hz with his backside facing the female (“peahen”). Next, during “train-rattling”, the male vibrates his tail and train at 25-28 Hz (mean 25.6 Hz) while facing toward the female at close range (1 to 1.5 m) (Fig 1A, S1 Movie), causing the train to shimmer iridescently and emit a prolonged “rattling” sound [18–20]. Train-rattling performance by peacocks is obligatory for mating success [18], and eye-tracking experiments have shown that both wing-shaking and train-rattling displays are effective at attracting and holding the peahen’s gaze [21]. Peahens also perform a tail-rattling display at 25-29 Hz in a variety of contexts [20], suggesting that feather vibrations might serve other communicative functions as well. Three studies have found that birds respond behaviorally to playbacks of low frequency sound generated by social displays: peafowl detect the infrasound (< 20 Hz) component of train-rattling and wing-shaking recordings [19]; male houbara bustards (*Chlamydotis undulata undulata*) respond to low frequency (40-54 Hz) boom vocalizations [22]; and male ruffed grouse (*Bonasa umbellus*) respond to 45 ± 6 Hz wing beating “drumming” displays [23]. Several other studies have also measured the behavioral response of birds to amplitude-modulated low repetition rate broad-band pulses, which are similar to the mechanical sounds associated with peacock train-rattling; the results of these studies showed that birds from three different families can detect such sounds with repetition rates < 40 Hz [24–26]. However, no studies have considered whether peafowl or other birds might detect such mechanical sounds via vibrotactile perception (i.e., sensing sound air particle velocity or airflow impulses via feather vibrations or deflections) as well as by hearing (sound pressure wave reception).

**Fig 1.**
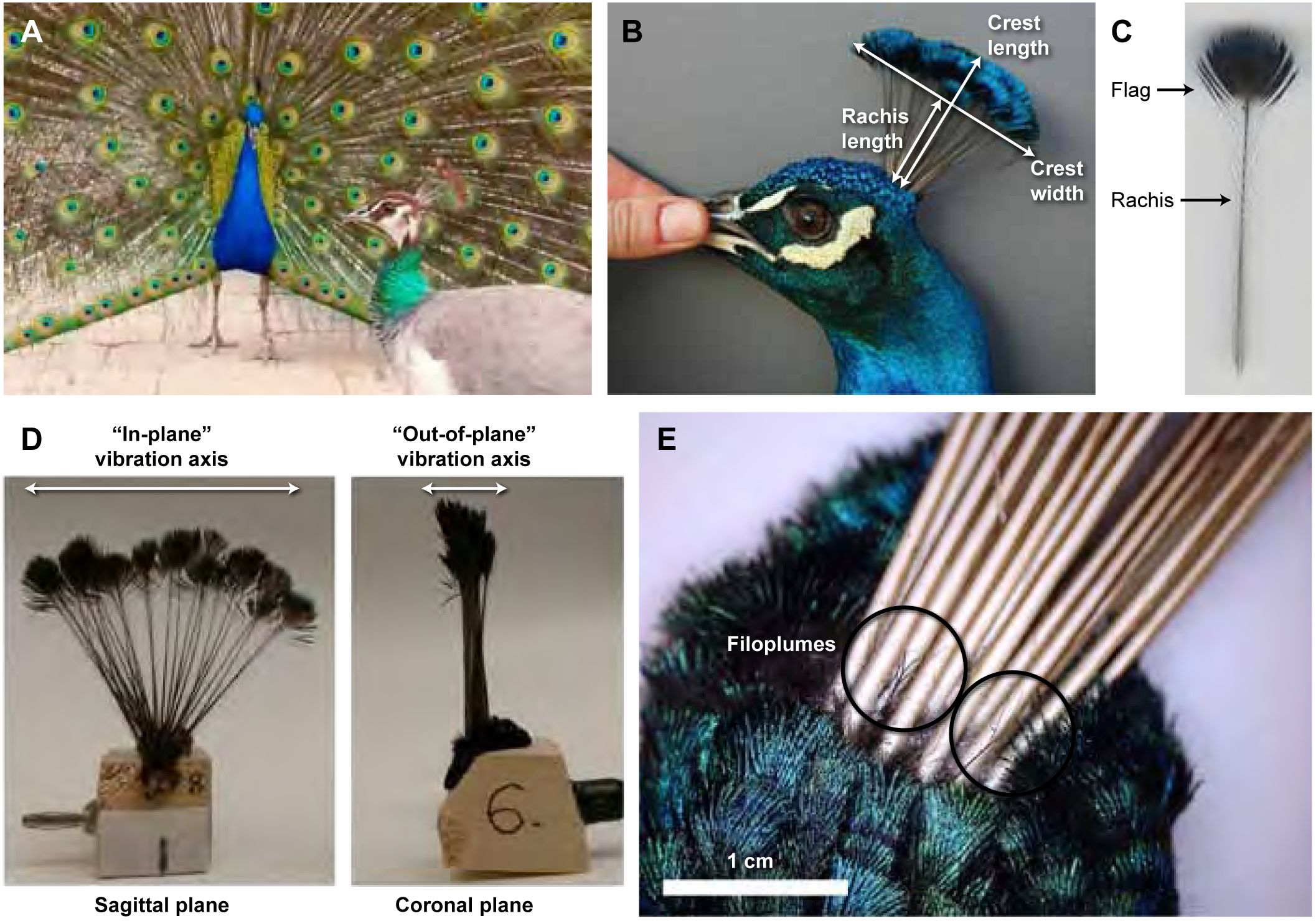
Morphology of peafowl crests and crest feathers. (A) A peahen (foreground) with the plane of her crest oriented towards the displaying peacock (background) as he performs train-rattling vibrations. (B) Both sexes have a crest with an inverted pendulum shape made up of between 20-31 feathers. This photo shows an adult male measured *in vivo*. (C) A single crest feather showing the pennaceous flag at the distal end. Note that only short, thin barbs are present on the relatively bare rachis (shaft) at the proximal end. (D) A whole crest sample mounted for the laboratory experiments. The two axes of vibrational motions (“in-plane” and “out-of-plane”) are indicated. (E) Mechanosensory filoplumes (circled) are located at the base of the peafowl crest feathers.

One possible means by which peafowl might sense sound by vibrotactile perception is the fan-like crest (Figs 1A,B), a planar array of feathers oriented in the sagittal plane that is found on the heads of both sexes [17]. Each crest feather has a spatulate “flag” of pennaceous vanes at the distal end and a long shaft that is mostly bare except for short, sparse barbs along its proximal end (Fig 1C). The pennaceous flag of peafowl crest feathers might couple to oscillations in surrounding air via drag forces induced by flow of the surrounding medium, as well as via forces exerted on the flag’s face by incident pressure waves.

We first consider the mechanical properties one would expect feathers to have to function effectively as mechanosensors, based on mechanoreception in other animals. Our predictions (P1-P4) are outlined in Box 1 below. A large body of research in mammals and arthropods has found that antennae and sensory hairs play important mechanosensory roles in sound detection; this function is also known to be influenced by their vibrational response and mechanical structures [27,28]. For example, in order for a feather crest to sense environmental airflows, it would need to bend sufficiently to activate mechanosensory nerve cells (P1-P4; Box 1), as has been shown for pigeon covert feathers [2], arthropod sensory hairs, pinniped whiskers, bat sensory hairs, fish lateral line organs [29], and rat whiskers [30,30]. Thus, one would expect crest feathers to be compliant enough to deflect when stimulated by salient airflow stimuli. Sound consist of oscillations of the surrounding medium in both pressure and particle velocity. Animals can detect pressure oscillations using ears and tympanal organs, whereas particle velocity oscillations can be detected in a variety of ways, including using sensory hairs and antennae that have mechanosensors at their bases [31,32]. Feathers of all types in birds of all orders have at their bases specialized short mechanosensitive feathers called filoplumes that couple to motions of their associated feather’s shaft [33,34]; thus, we expect this to also be true for crest feathers (prediction P1; Box 1). Like many sensory hairs and antennae, the plumose structure of feathers enables effective mechanical coupling to air motions via drag forces [2,35] and elongated, tapering shafts well-suited for bending and transmitting force to an enervated base. Both contour feathers and filoplumes have been shown empirically to detect bending and vibrations via mechanoreceptive Herbst corpuscles at their bases [36,2,3,37].

Because the peafowl’s region of most acute vision is oriented laterally [38], when a peahen gazes at a displaying male, the maximum area of her crest feathers also points toward the peacock’s moving feathers (Fig 1A). This results in an optimal orientation for intercepting airborne vibrations generated by display behaviors; these medium oscillations should tend to drive feather crests to oscillate in the “out-of-plane” orientation (i.e., normal to the plane of the crest as shown in Fig 1D). The displaying peacock shakes its body laterally during such behaviors, so any corresponding vibrations of its own crest should also occur in the out-of-plane direction. The design of the crest also provides a mechanical advantage, enabling it to transmit a magnified version of the forces applied to its distal end to its base, because the flag is wider than the tapered base and because each feather shaft acts as a lever arm coupling the flag to the base.

#### Box 1. Predicted properties of feathers that detect airborne stimuli

P1. Coupled to mechanosensory structures. *(Morphology; Figs 1-2)*

P2. Frequency-tuned to stimuli with well-defined frequencies. *(Vibrational dynamics measurements; Fig 3)*

P3. Damped at the right level to allow detection of airflow impulse rate. *(Force impulse experiments; Fig 4)*

P4. Responsive to experimentally-simulated social stimuli. *(Audio playback experiments, Simulated wing-shaking experiments; Figs 5-6)*

Another important consideration is the frequency-tuning between sensory structures and their stimuli (prediction P2; Box 1), as this can provide several advantages including filtering out background noise and other irrelevant stimuli [31]. In fact, a variety of arthropods use receivers with a response that is frequency-matched to the stimulus source, including antennae and sensory hairs used by various insect species to detect wingbeat signals of near-by conspecifics, trichobotheria used by some arachnids to sense prey wingbeats [39–43], and frequency-tuned eardrums used by cicadas and crickets to detect conspecific songs [44,45]. Such mechanical frequency tuning can be accomplished readily via resonance, the phenomenon whereby an object responds with maximum amplitude to a driving force that oscillates near one of its natural frequencies of vibration [31]. Resonant frequency matching enables a mechanoreceptor to respond with optimal sensitivity to low amplitude airborne stimuli, at the expense of frequency discrimination (by contrast, human eardrums and microphones have broadly-tuned resonance responses that allow efficient detection of natural stimuli over a wide range of frequencies).

Therefore, given that peafowl displays take place at well-defined shaking frequencies, we predict that their feather crests might have a resonant frequency response with a peak and width matched to the display, as found for the arthropods cited above. If this frequency tuning is indeed present, the crest might enable peafowl to detect airborne stimuli generated by a conspecific individual’s shaking motions by undergoing sympathetic vibrations (i.e., undergoing oscillations driven by coupling through drag forces to air particle velocity oscillations). Feather crests optimized to vibrate at the shaking frequency could also provide proprioceptive feedback to the individual performing the display [46,47,31].

Some arthropods have the additional ability to use mechanosensory hairs to sense separate airflow pulses generated by abrupt, repetitive motions. The resulting impulsive forces cause the mechanosensors to oscillate only briefly at their natural frequency before their motion is damped out. For example, the cerci sensory hairs of female African crickets (*Phaeophilacris spectrum*) function in this way to detect air vortices produced by males performing wing flicks [48–50]. These motions are similar to those performed during peacock wing-shaking displays. Detecting airflow impulses via transient oscillations requires the right level of damping and natural frequency to allow a high amplitude response while also enabling detection of the airflow impulse repetition rate (prediction P3). Consequently, feather crests would also need this specific combination of vibrational response parameters in order to respond efficiently to impulsive airflows generated by wing motions.

In other animals, vibrotactile sensors detect sound particle velocity oscillations in the acoustic near-field, a region close enough to the source that particle velocity can couple efficiently to mechanoreceptors via drag forces [31]. For example, in arthropods, many species use filiform hairs to detect near-field particle velocity for predator or prey detection and for intraspecies signaling [51–53]. Near-field communication has been studied in a wide variety of invertebrate terrestrial taxa [54,51] and in fish [55]. By contrast, it is often assumed that only the acoustic far-field is relevant for sound reception by birds. In the far-field, sound predominantly consists of pressure waves detectable by vertebrate ears, insect tympanal organs and similar receptors. Because the particle velocity magnitude falls off more rapidly with distance than the pressure wave component, particle velocity stimuli are greater than those due to pressure waves only for distances R < 0.16 to 0.22 λ (where λ = wavelength) for monopole sources (e.g., loudspeakers) and dipole sources (e.g., moving wings, tails and trains), respectively [56,31,57]. Consequently, this wavelength-dependent distance is often used to distinguish the acoustic far-and near-fields. However, the relevant criterion for efficient mechanosensation is the *absolute magnitude* of particle velocity, not the *relative value* of particle velocity compared to the pressure wave [58]. Thus, the regime relevant for vibrotactile sensing is the flow (reactive) near-field: the region near the sound source where the particle velocity has its greatest magnitude because the air acts as a layer of effectively incompressible fluid that moves with the source [32,59]. The extent of the flow near-field depends on source size, A, not wavelength (see e.g., Fig 2 in [59]). For R ≤ 0.16 A (the “very near-field”), the particle velocity is approximately constant. As R increases, particle velocity becomes negligible for mechanosensing at approximately R ≈ A [31,59,35]. The lateral extent of the flow near-field also depends on A. In addition, the overall magnitude of the particle velocity increases as A^2^ for a monopole and A^3^ for a dipole. In summary, increasing source size, A, increases the spatial extent of the flow near-field regime in which mechanoreception can take place, as well as the magnitude of particle velocity stimuli [56].

**Fig 2.**
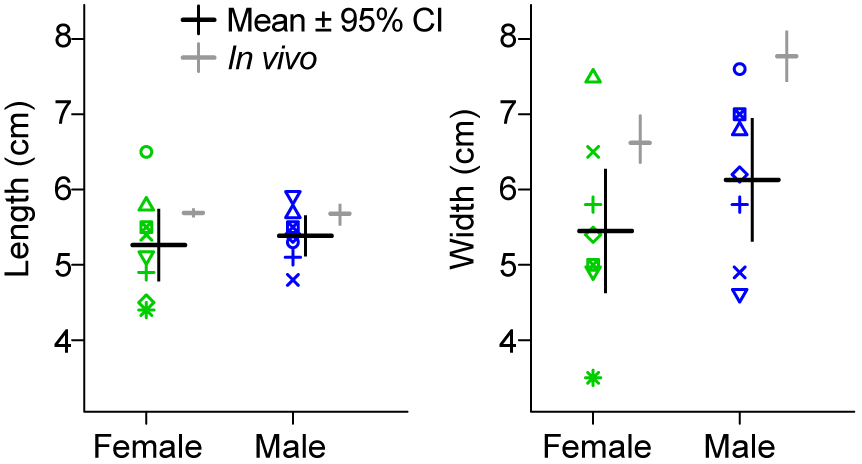
Length and width of the whole crest samples as compared to live peafowl crests. Crests (n = 8 female, n = 7 male) measured *in vivo* (means shown to the right of each data column) had similar morphology to the dried samples, except that the crests on live birds tended to be wider. Dried sample dimensions were measured to the nearest 0.1 cm. Each crest sample is indicated by a unique symbol-color combination consistent with other figures (see S4 Table for details).

During the peacock’s display, typical female-male distances, R = 1.0 to 1.5 m, are equal to the typical peacock train radius, which plays the role of source size A. Female therefore experience train-rattling sound in the acoustic flow near-field [18,60], satisfying a prerequisite for vibrotactile sensing. The criterion R = 1 to 1.5 m < 0.22 λ corresponds to frequencies < 50 to 75 Hz at this distance; therefore, the mechanical sound generated by the peacock’s display has appreciable pressure wave magnitude as well for most frequencies in the human audible range. Spectrograms from earlier studies of peacock train-rattling indicate that these mechanical sounds consist of broadband impulsive rattles with spectral density primarily in the human audible range, emitted at a repetition rate of approximately 26 Hz; they are neither low frequency pure tones, nor are they sound with spectral density predominantly in the low frequency or infrasound regime [19,20]. As a result, a consideration of the sound fields of peacock train-rattling displays indicates that rattling sounds might be detectable as particle velocity or pressure wave stimuli, or both, at typical display distances.

In this study, we compare the mechanical properties of peafowl crests with those predicted for mechanosensation (predictions P1-3; Box 1), and we furthermore test whether stimuli from peacock displays induce a vibrational response in the crest (prediction P4). We also wished to determine whether any agreement between social display frequencies and crest resonant frequencies was generic or specific to this system. Therefore, we sought to understand how the peafowl crest’s resonant properties relate to those of other types of peafowl feathers as well as crest feathers from other species. This work was designed to serve as a first step to determine whether the head crests of birds might serve a variety of mechanosensory functions, including the detection of body self-motion and various airborne stimuli.

## Materials and methods

### Morphology

A total of n = 7 male and n = 8 female Indian peafowl (*Pavo cristatus* Linnaeus 1758) head crests with the feathers still mounted in skin were obtained from Moonlight Feather (Ventura, CA USA) and Antebellum Anne (Pell City, AL USA); other peacock samples and crest feathers from four other bird species were obtained from Moonlight Feather (Ventura, CA USA), Assiniboine Park Zoo (Winnipeg, Manitoba, Canada) and Siskiyou Aviary (Ashland, OR USA) (see S1 Table and S1 Fig for details).

Motivated by reports that mechanosensitive auklet crest feathers are filoplumes [14] and that filoplumes from diverse species spanning several orders might function to detect disturbances of the surrounding feathers [61–64], we used microscopy to determine whether peafowl crest feathers either are themselves filoplumes or have filoplumes at their bases. A Digital Microscope Pro (Celestron, Torrance, CA USA) was used to examine the base of peafowl crest feathers to determine whether filoplumes were present, using the structural criteria employed in previous studies of this feather type (i.e., short feathers with a long, bare shaft with a tuft of short barbs on the distal end, located near the base of a longer feather but not growing from the same follicle [61,33,62,64,63,65]; see micrographs in [33,36,65]. Crest length and width measurements were made by hand and from digital photographs of the crest samples and high-resolution scans (0.02 mm/pixel) of single feathers with a ruler included in the sample plane. We used these measurements to compare the morphology of dried crest samples with that found for crests on live peafowl in a previous study [17], including length, width and number of feathers. Because some peafowl, especially females, have non-uniform crest feather lengths [17], we also measured the lengths of individual feathers within the dried crest samples to compare with the previous study. If the crest feathers were closely clustered, the attached skin was first softened in water and the crest was spread to approximate its natural configuration.

Following earlier studies of feather vibrational properties [20,66,67], we mounted crest feathers by gluing the crest skin to a rigid sample holder (a 2.5 cm cube of balsa wood) (Fig 1D). This method is justified because the resonant frequency of a flexible shaft secured at one end by a stiff clamp is not expected to be affected by the clamp’s mechanical properties [31]. In an earlier study, we had verified that this is true for peafowl tail and train feathers [20]: i.e., there was minimal shift (< a few percent) in feather resonant frequencies in the frequency range considered in this study when samples were mounted on rigid wooden blocks vs. embedded in a compliant gel to mimic the shaft’s native soft tissue environment. Further supporting this method, we found good agreement between train-rattling display shaking frequencies and the value predicted from a model of the peacock tail’s resonant frequency based on laboratory measurements [20]; in addition, an earlier study of manakin feather resonance that used similar mounting methods found good agreement between the frequencies of feather vibrational resonance measured in the laboratory and sonations recorded in the field [66].

Because interactions between feathers can influence their resonant frequency and damping [66,68], we compared the biomechanics of whole crests to that of isolated crest feathers. To study individual, isolated crest feathers, we removed all but three to five feathers (on the outer edges and in the middle) from two male crests and one female crest and analyzed the characteristics of those remaining feathers. Note that because this procedure was necessarily destructive, it precluded any further whole-crest analyses on those samples, we limited it to only the three crests. Isolated body and crest feathers were inserted into a close-fitting hole in the wood base using polyvinyl acetate glue. For measurements on other peafowl feathers and crest feathers from other species, all but two feather samples in S1 Table were inserted up to the top of their calamus (the part of the shaft inserted into the skin). The Victoria crowned pigeon crest feathers had been trimmed just above the calamus, so they were mounted such that 3 mm of the exposed feather shaft (3% of its length) was inserted into the wooden base.

To ensure further that the feathers had the same mechanical properties as those found on live birds, we stored and tested all samples using environmental conditions similar to those measured during the peafowl behaviors of interest. Because earlier research on feather keratin indicated that water content can affect its elastic modulus [69], all crest samples were stored and all laboratory measurements were taken at 21.1 C° (range: 20.8-21.5°) and 74.8% relative humidity (range 72.3-77.7%). For comparison, peacock train-rattling display frequencies were measured in the field at a median temperature of 19.4°C and a median relative humidity of 60.7% [20], with over half of the displays occurring within ±2.2°C and ±14% of the average laboratory temperature and relative humidity, respectively. Moreover, a re-analysis of previous published data on 35 peacock displays performed by 12 males in the field [20] shows that there is no significant association between display vibration frequency and relative humidity (p > 0.45) when accounting for the date and time of the displays. This analysis and the associated data are provided in the data repository for this study [70]. As an additional check, we also measured the audio playback response with crest samples held at 35% relative humidity and 22 C° for 1 min to 10 min and found no measurable change in the natural frequency over this time.

### Ethics statement

All research procedures were approved by the Haverford College Institutional Animal Care and Use Committee (protocol #sak_050916).

### Vibrational dynamics measurements

To determine the vibrational resonant frequency of each feather sample, and its relationship to possible driving mechanisms during displays, we applied a sinusoidal force to the sample while measuring its resulting vibrational amplitude as a function of the driving force’s frequency. The system’s vibrational response (transfer function) is then computed as the ratio of the sample’s amplitude of response to the driving stimulus magnitude [71,66,67]. For these measurements we mounted each feather sample on a model SF-9324 mechanical shaker (Pasco Scientific, Roseville, CA, USA) driven by an Agilent 33120A function generator (Agilent Technologies, Wilmington, DE, USA) (S1 Fig). This apparatus applied sinusoidal forces with a linearly varying frequency (“frequency sweeps”) while high-speed video was used to measure the amplitude and frequency of vibration of both the driving mechanism and the feather sample (details on the frequency sweep parameters and video methods are discussed below). The driving force was applied in two orthogonal directions (Fig 1D): 1) “out-of-plane” (oriented normal to the plane of the crest), corresponding to the geometry when a peafowl views a display with its laterally-oriented visual field, or drives its own crest into vibrations by performing a train-or tail-rattling display [20]; and 2) “in-plane” (oriented parallel to the plane of the crest, in the posterior-anterior axis of the head), corresponding to the geometry when the front of the head is oriented towards the display.

The resulting vibrational response spectra of the crests were measured using three linear frequency sweeps. One of these sweeps used the frequency range (0-80 Hz) to include all peaks in the spectral response found for peacock tail and trail feathers in an earlier study [20]; the rate of frequency increase (1.33 Hz/s) was chosen to be less than the values measured at the start of peafowl displays in the same study. These conditions were used to test the vibrational response in the out-of-plane direction for each of the 15 peafowl crests, as well as three of the crests that had been trimmed down to have only three to five isolated crest feathers remaining (n = 3 trials for each sample); this allowed us to compare the vibrational response of intact crests with that of isolated crest feathers. We ran the following additional trials to make sure that this combination of frequency range and sweep rate above did not miss any spectra peaks or affect the shapes of the resonant peaks: six crests out-of-plane at 10-120 Hz (1.83 Hz/s; n = 18 trials), six crests out-of-plane at 0-15 Hz (0.25 Hz/s, n = 6 trials), five crests in-plane at the 0-80 Hz range (1.33 Hz/s; n = 14 trials), and two crests in-plane at 10-120 Hz (1.83 Hz/s; n = 2 trials). For one crest, we determined that varying the amplitude of shaking by a factor of four resulted in the same resonant response within measurement error.

We also measured the resonant vibrational response of crest feathers for several other types of short peacock feathers (three lengths of peacock mantle feathers, the shortest length of train eyespot feather, and four different body contour feathers), and for one or more crest feathers from four other bird species: two additional species from order Galliformes, the Himalayan monal (*Lophophorus impejanus*) and the golden pheasant (*Chrysolophus pictus*); the Victoria crowned pigeon (*Goura Victoria*) from the order Columbiformes, and the yellow-crested cockatoo (*Cacatua sulphurea*) from the order Psittaciformes; see S1 Table and S1 Fig for details. The resonant response for each of these feathers was measured for driving forces in the out-of-plane direction for n = 3 trials for each feather at each of two frequency sweep rates (2.0 Hz/s, 0 to 120 Hz; 1.33 Hz/s, 0 to 80 Hz); the Himalayan monal sample was also studied using a sweep rate of 0.5 Hz/s over 0 to 30 Hz because it had a lower frequency response.

### Video analysis

We recorded feather vibrational motions using high-speed video filmed with a GoPro Hero 4 Black Edition camera (720 x 1280 pixels; 240 frames s^-1^; GoPro, San Mateo, CA, USA). Similar video and imaging-based methods have been used to measure resonance in whiskers [71–74], insect antennae [75], feathers [76] and human-made structures [77,78]. Image and data analysis were performed using custom programs based on the MATLAB 2015a Machine Vision, Signal Processing and Curve Fitting toolboxes (MathWorks, Natick, MA, USA); the MATLAB scripts to reproduce this analysis are available with the data repository for this study [70]. The Nyquist frequency, which gives the upper bound on measurable frequencies [79], was 120 Hz (half the frame capture rate) (> 4× typical biological vibration frequencies used during peafowl displays). Images were first corrected for lens distortion using the MATLAB Camera Calibration tool. All feather motions analyzed were in the plane of the image, and thus did not require correction for perspective [80]. To analyze feather motion, we first used auto-contrast enhancement and thresholding to track the mean position of the crest feather flags and the shaker mount, and then computed the spectrogram of each object’s tracked position during the frequency sweep using a Hanning filter. This yielded the magnitude of the fast Fourier transform (FFT) at each vibrational drive frequency, *f_d_*, measured for motion of the crest flag, *A*, and that of the sample holder, *A_d_*, which provides the driving force. Mechanical shakers have a frequency response that necessarily rolls off in amplitude at the low frequencies considered here due to fundamental physical principles [79]; see, e.g. inset to Fig 1A in [81]. To account for frequency-dependent variation in the driving force, we then divided the sample’s magnitude, *A*, by the shaker drive magnitude, *A_d_*, at each drive frequency, *f_d_*, and smoothed the ratio over a 1.3 Hz window using a cubic Savitzky-Golay filter to give the drive transfer function, *H(f_d_)* = *A*/*A_d_* [71,66]. Nonlinear least squares fitting using Origin 8.6 (Originlab, Northampton MA USA) was used to fit each peak in the transfer function, to a Lorentzian spectral response:

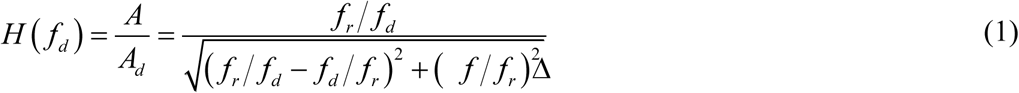

where the fit parameters are *f_r_*, the resonant frequency, and Δ*f*, the full-width-half-maximum of the spectral power; this yielded mean and s.e.m. estimates for the fit parameters as well as the quality factor, *Q* = *f_r_* /Δ*f*, a measure of how sharply the transfer function is peaked about the resonant frequency [79].

### Force impulse experiments

A second standard method for determining the vibrational response of a system involves applying a transient impulsive force and then measuring the system’s subsequent vibrational response to measure its natural frequency of vibration, *f_o_*, and the exponential decay in time of its vibrational amplitude [71,82,72,75,76]; this is analogous to striking a bell and recording how it rings at a well-defined frequency as its sound intensity decays in time. This method is also relevant for determining the response of the crest to impulsive airflows due to each flap of the wing during wing-shaking. Thirdly, it also serves as a check on the validity of the vibrational response frequency sweep methods described above, because the natural and resonant frequency should be related as [79]:

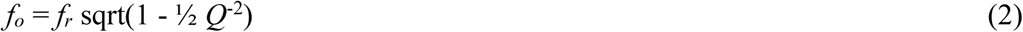

This prediction can be tested by comparing the natural frequency measured directly from the force impulse method with the value computed from the vibrational response’s transfer function using Eq 1 and 2.

To determine the peafowl crests’ response to impulsive airflows, we impacted crests with single air ring vortices and measured the resulting motions on video. A Zero Blaster vortex gun (Zero Toys, Concord, MA, USA) was used to generate single air vortex rings of artificial fog (2-4 cm in diameter, 1 cm diameter cross-section, speed 1.8 m/s [95% CI 1.7, 2.0 m/s, range 1.5 - 2.1 m/s]), aimed so as to impact whole crests (n = 2 peacock and 1 peahen) in the out-of-plane orientation. The motion of crest feathers struck by the vortices was measured by tracking the crest position on high-speed video when an intact vortex impacted the crest oriented with its widest cross-section facing the source at 0.5 m from the point of creation. As explained above, because we expected such impulses to result in the crest feathers oscillating at their natural frequency, this provided an additional check on our resonant frequency values. This also provides a model for understanding how crest feathers would respond to impulsive airflows generated by other sources (e.g., displays, wind, etc.).

### Audio playback experiments and analysis

To determine if peafowl crests can vibrate detectably due to peacock train-rattling, we filmed high-speed video of peahen crest samples placed in the flow near-field of a loudspeaker playing back train-rattling sounds. Note that because peacock train-rattling consists of broad-band rattles at low repetition rates, not pure tones, we used audio equipment rated for frequencies 20 Hz to 20 kHz rather than equipment designed for infrasound, similar to how one would treat the sound of hands clapping or birds calling at a repetition rate of a few Hz. To generate audio playback sequences, we used audio field recordings (24-bit, 44.1 kHz, no filtering) of peacock train-rattling displays made using a PMD661 recorder (±1 dB: 20 Hz to 24 kHz; Marantz, New York, NY, USA) and a ME-62 omnidirectional microphone (±2.5 dB: 20 Hz to 20 kHz; Sennheiser, Wedemark, Germany), as described in a previous study [20]. Three playback sequences were used (each using sound from a different peacock), with mean rattle repetition rates of 26.7 ± 0.5 Hz; 25.3 ± 0.5 Hz; and 24.6 ± 0.5 Hz. Recordings of train-rattling in the field indicated that rattles are in-phase (i.e., temporally coherent) over bouts approximately 1.2 s duration that are repeated for several minutes during displays [18,20]. We spliced together bouts with an integer number of rattling periods to form a longer audio playback file with a total duration of approximately 5 min.

All sound files were played back on a Lenovo Thinkpad T460S computer connected to a 402-VLZ4 mixer (Mackie; preamplifier; < 0.0007% distortion 20 Hz to 50 kHz) and a ROKIT 10-3 G3 10" powered studio monitor (KRK Systems, Fort Lauderdale, Florida, USA; ± 2.5 dB over to 40 Hz to 20kHz, –10 dB at 25 Hz relative to ≥ 40 Hz) with a 25.4 cm diameter subwoofer (A = 12.7 cm). Following [83], we examined re-recordings of the playback stimuli made with the same microphone and recorder used for the original field audio recordings, and found that the resulting waveforms and spectrograms (e.g., S2 Fig) had the same temporal features (“rattle” notes) as the original field recordings of train-rattling (e.g., Fig 4A in [20]).

As a control, we played back a Gaussian white noise file generated using MATLAB’s imnoise command (5 min. duration, 24-bit, 44100 Hz, FFT amplitude flat from < 1 Hz to 22,000 Hz computed using a rectangular window to preserve Fourier amplitudes). The white noise playback assessed whether the crest samples could be driven to vibrate measurably by a broadband signal similar to the train-rattles, but lacking the low-frequency amplitude modulation of the train-rattling recording at the “rattle” repetition rate. The root-mean-squared (rms) amplitudes of all playback recordings and the white noise control were scaled to the same value while also ensuring that no clipping occurred at high amplitude.

For playback experiments, the preamplifier volume controls of the mixer were adjusted so that the mean playback SPL was 88 ± 1 dB at 3 m as measured by a Type 2 model R8050 sound level meter (accuracy ±1.4 dB, C-weighting, 30-100 dB, slow 1.0 s setting; Reed Instruments, Wilmington, NC USA). For comparison, previously-reported values for peacock train-rattling mechanical sounds corrected for background noise were given as 67 to 77 dB at R = 3 m (unweighted SPL) [19], similar to audible bird wingbeat SPL summarized in S2 Table. Based on the frequency response specifications for our microphone and playback system, we estimated their combined response at 26 Hz to be reduced by 12.5 dB compared to the audible range; the increase in playback SPL relative to the reported values accommodates for this attenuation. During playbacks, the background noise with no audio was 58 ± 1 dB SPL.

Another consequence of the broadband nature of train-rattling is that rapid intensity variations due to interference at small R (the “interference near-field”) should not be relevant for this display, because this effect depends on the superposition of sound waves with a well-defined frequency emitted by different parts of an extended source. The predicted interference near-field regime is *R* < 2*π A*^2^/*λ* = 0.7 cm for 26 Hz [32]. Consistent with this expectation, we found no variation due to interference when we measured SPL at nine different positions across the subwoofer speaker between the center and edges, at distances perpendicular to the speaker between 12.7 cm to 0.5 m.

Female crest samples (n = 3; crests 7, 8, 15) with resonant responses determined in the vibrational dynamics trials were mounted on a tripod at a distance R = A = 12.7 cm away from the subwoofer speaker face to give optimal exposure to the flow near-field (S2 Fig). Vibrational motion of the samples was measured for three separate trials per crest and per recording using high-speed video (reducing speaker volume to zero and waiting > 5 s in between trials), and the crest motions were tracked and analyzed from video using the methods described above. The duration of train-rattling bouts gave an FFT frequency resolution of ± 0.50 Hz for vibrational response analysis. To minimize direct mechanical coupling via the substrate, the crest samples and speaker were mounted on Sorbothane™ vibration-isolation pads. Because peacocks often display near the edges of thick vegetation, next to natural ground slopes, or next to hard walls [84; personal observation], anechoic conditions are not required for effective courtship displays or for simulating their mechanical sounds. However, we still chose to minimize reverberations by surrounding the experiment with acoustic tiles and sound absorbing sheets (Audimute, Cleveland, OH USA; audible sound reduction rating: SAA 0.68, NRC 0.65), resulting in an SPL decrease of 5 dB when distance was doubled for R ≥ 0.25 m. This decrease in SPL is intermediate between the free-field value of 6 dB and a typical reflective room value of 3 dB [85]. We also performed negative controls to ensure that reverberations and substrate vibrations did not drive crest vibrations. This was accomplished by inserting a foam tile between the crest samples and the speaker to block particle velocity oscillations and attenuate directional sound pressure waves from the speaker. Thus, any crest vibrations measured during the controls would be due to substrate vibrations, reverberations, transmitted sound pressure waves, and/or other environmental sources.

### Simulated wing-shaking experiments

High-speed videos from a previous study were used to determine the frequency and amplitude of wing motions during the peacock’s wing-shaking display [20]. We used four videos filmed with the wingtip motion closely aligned with the image plane (see S1 Movie) that also showed tail feathers with known lengths. The amplitude of wing-shaking motion was defined by the mean diameter of motion circumscribed by the tips of the partly-unfurled wings during this display, which we estimated to be 7.6 cm on average (range 5.5 to 10 cm). To simulate the wing motions observed in displaying peacocks and the resulting air motions, we used a robotic mechanism that caused an entire peacock wing to flap with the wing plane held in a fixed vertical orientation while the wingtip circumscribed a circle (S1 Movie and S3 Fig). The peacock wing was mounted on a carbon fiber rod using a balsa wood base that was attached to the wing via adhesive at the shoulder; this rod pivoted about a clevis joint, which allowed the wing axis to move in a vertical circle while the wingspan remained in the vertical plane. At the end opposite the wing, the rod was attached to a circular crank by a universal joint. The crank and attached wing assembly was driven at 4.95 ± 0.05 Hz by a DC motor. To account for the fact that actual wing-shaking involves motion of two wings toward each other, which presumably displaces more air than a single wing, this apparatus used a single flapping wing moving in a slightly larger diameter (14 cm) circle at the wingtips.

To determine how wing-shaking influences the crest of an observing bird, we first determined the location of maximal airflow speed during robotic wing-shaking. Airflow speeds were measured by a model 405i Wireless Hot-wire Anemometer (Testo, Sparta, NJ, USA) oriented with its sensor facing in the same direction as the crest samples; this device has a resolution of 0.01 m/s, accuracy of 0.1 m/s, measurement rate of 1 Hz, and equilibration time of approximately 5 s. To define the airflow pattern around the flapping wing, air speed was sampled at every point on a 5 cm grid, 5-7 times per location. Based on these results, we determined the angle at which to position the crest sample. S3 Fig shows how three peahen feather crest samples (Crests 08, 12, and 13) were positioned using a tripod at the vertical midline of the wing located at various distances from the wing-tips. The resulting motion of the crests was then filmed using high-speed video as described above in “Video analysis” to quantify the vibrational response of the three peahen crests. To verify that substrate vibrations did not drive the crest motion, we also performed a control by inserting a 3 x 4 ft foamboard in between the crest and wing to block the airflow from the wing motion; this reduced the root-mean-squared crest motion to 14% of its value with wing motion-induced airflow present. For comparison with the wing-shaking frequency during displays, flapping frequencies during ascending and level flight were also measured for 9 peacocks from 6 online videos (S3 Table).

### Force measurements

Peacock feather keratin, like other biopolymers, can have a nonlinear elastic response to external stresses [86]. Because the stimuli in the mechanical shaker, audio playback and wing-shaking experiments each exerted different forces and these forces may have had greater magnitudes than those encountered in the field, we wanted to understand how to extrapolate from our laboratory experiments to a lower force regime that is potentially more biologically relevant. Consequently, we measured the elastic mechanical response of peafowl crests to an external bending force applied to the flags of the crest. We studied the static mechanical response of peafowl crests in the single cantilever bending geometry by measuring the relationship between flag displacement and restoring force of the crest in the out-of-plane orientation (Fig 1D). Force measurements were made using a Model DFS-BTA force sensor (accuracy ± 0.01 N) connected to a LabQuest2 datalogger (Vernier Software & Technology, Beaverton, OR, USA), which was calibrated using known masses. The force sensor was attached to a thin rectangular plastic blade oriented in the horizontal plane. The edge of the blade was pressed against the midpoint of the flags of the vertically oriented crest to measure the restoring force exerted by the bent crests. The crests were mounted on a micrometer that moved them toward the force sensor and enabled measurement of crest displacement relative to the location at which the crest flag first deformed and the restoring force first became non-zero within measurement error. These measurements were performed for three trials each for three male and three female crest samples. The resulting force vs displacement data were fit to a linear force-displacement model to determine the linearity of elastic bending deformations. This also gave a value of the bending spring constant, k.

### Statistical analysis

All measurements and fitted values are reported as means [95% confidence interval, defined as 1.96 × s.e.m. for normally distributed data], unless noted otherwise. To analyze sources of variation in whole crest *f_r_* and *Q*, we fit Gaussian linear mixed-effects models with a random intercept of crest ID to account for repeated measures of each bird’s crest using the nlme 3.1-131 package [87] in R 3.3.3 [88]. We first verified that trial order and frequency sweep rate, two aspects of the experimental design, did not have significant effects on either *f_r_* or *Q* (all p > 0.28). The next step was to evaluate the potential effects of morphological traits that could influence crest resonance (some of which could be weakly correlated in a much larger study; see [17]). Because our sample size was only 15 crests, but we had five morphological traits, our statistical power was only sufficient to consider models with only one morphological trait predictor at a time: length, width, number of feathers, percent of unaligned feathers, and percent of short feathers. These morphological traits were fitted as fixed effect predictors. All models also included fixed effects of sex as well as the vibration orientation (either in-plane, or out-of-plane). We used AICc to select the best-fit model [89] and evaluated significance of the fixed effects using Wald tests. We report *R*^2^_LMM(m)_ as a measure of the total variance explained by the fixed effects [90,89]. We also used the variance components of the best-fit model to calculate the adjusted repeatability, defined as the variance attributed to differences among crests after adjusting for variation explained by the fixed effects [91]. Inspection of the data and model residuals revealed that variance in *f_r_* differed among crests, so when modelling *f_r_*, we fit a heteroskedastic model that had its standard errors adjusted to account for the appropriate within-group error variance, by using the varIdent option in the weights argument in nlme [87].

## Results

### Morphology

A microscopic examination of peafowl crest feathers reveals that their shafts have associated feathers with the structure of filoplumes at the base (Fig 1E) that agree in location and morphology with those shown in micrographs of filoplumes cited earlier in the Methods. These were structurally distinct from immature crest feathers, which also retained a sheath until they had grown to a length much greater than that of the filoplumes.

The average lengths of the whole crest samples used in this study were 5.3 [4.8, 5.7] cm for 8 female crests, and 5.4 [5.1, 5.7] cm for 7 male crests. Fig 2 shows that this range of crest lengths agrees with that of live peafowl, indicating that the crest samples used in these experiments were fully grown [17]. The average widths of the whole crest samples were 5.5 [4.6, 6.3] cm for the female crests, and 6.1 [5.3, 6.9] cm for the male crests. These width values were approximately 20% (female) to 27% (male) smaller than those found on live birds (Fig 2). This difference could be due to the crest ornament being spread 1-2 cm more in the sagittal plane by muscle action in the live bird, as observed for erectile crest plumage in many other species [13], in addition to the effect of skin drying.

All 7 of the male crest samples had feathers of uniform length, defined as ±8% of the mean fully-grown crest feather length. This is also typically observed *in vivo*, where 72% of male *P. cristatus* crests studied in [17] had feathers of uniform length. In contrast, the majority (6/8, or 75%) of the female crest samples had non-uniform feather lengths (using the same definition above), which was again similar to the previous *in vivo* study, where 77% of females had non-uniform crest feather lengths [17]. On average, the dried female crests had 7.0% [2.1, 11.8] of their feathers shorter than the mean fully-grown crest feather length. Eight out of the 15 crest samples had all feathers oriented in the same plane within ±5°; five of the crests had 7-11% of the feathers unaligned, and two male crests had 22% and 50% unaligned feathers, respectively.

We also studied the morphology of individual peafowl crest feathers to understand their unusual structure (Fig 1C). The average rachis tapered evenly over its 39.90 [38.89, 40.91] mm length and had a mass of 5.1 [4.8, 5.3] mg, and the plume (or flag) added another 2.50 [0.87, 4.06] mg. Unlike the fully formed barbs in the pennaceous flag, the lower barbs were short (4.1 [3.0, 5.2] mm) and lacked barbules altogether.

### Vibrational dynamics measurements

The vibrational drive transfer functions of peafowl crests had either a single dominant fundamental peak, or in a few cases, a cluster of two to three peaks in a narrowly-defined frequency range, with no evidence that other modes of vibration caused detectable motions of the pennaceous flags. The functional form of each main spectral peak agreed well with the Lorentzian (mean adjusted-*R*^2^ = 0.97; range [0.91, 0.998]) (Eq 1) predicted for a cantilever [92], indicating that the system responded in the linear regime for our shaker amplitudes and frequency sweep rates (Fig 3A). The value of *f_r_* ± Δ*f* /2 defines the approximate range of drive frequencies over which power is efficiently coupled into the oscillator. Fig 3B shows that shaking frequencies measured in the field for displaying male and female peafowl [20] lay within *f_r_* ± Δ*f*/2 of the crest resonant frequency for both sexes (n = 8 female crests and 7 male crests). When the shaking force was oriented out-of-plane, the mean crest resonant frequency, *f_r_*, was 28.1 [28.0, 28.1] Hz for female and 26.3 [25.9, 26.6] Hz for male crests, respectively. The mean Δ*f* values were 6.2 [4.4,8.0] Hz (females) and 4.3 [4.2, 4.4] Hz (males).

**Fig 3.**
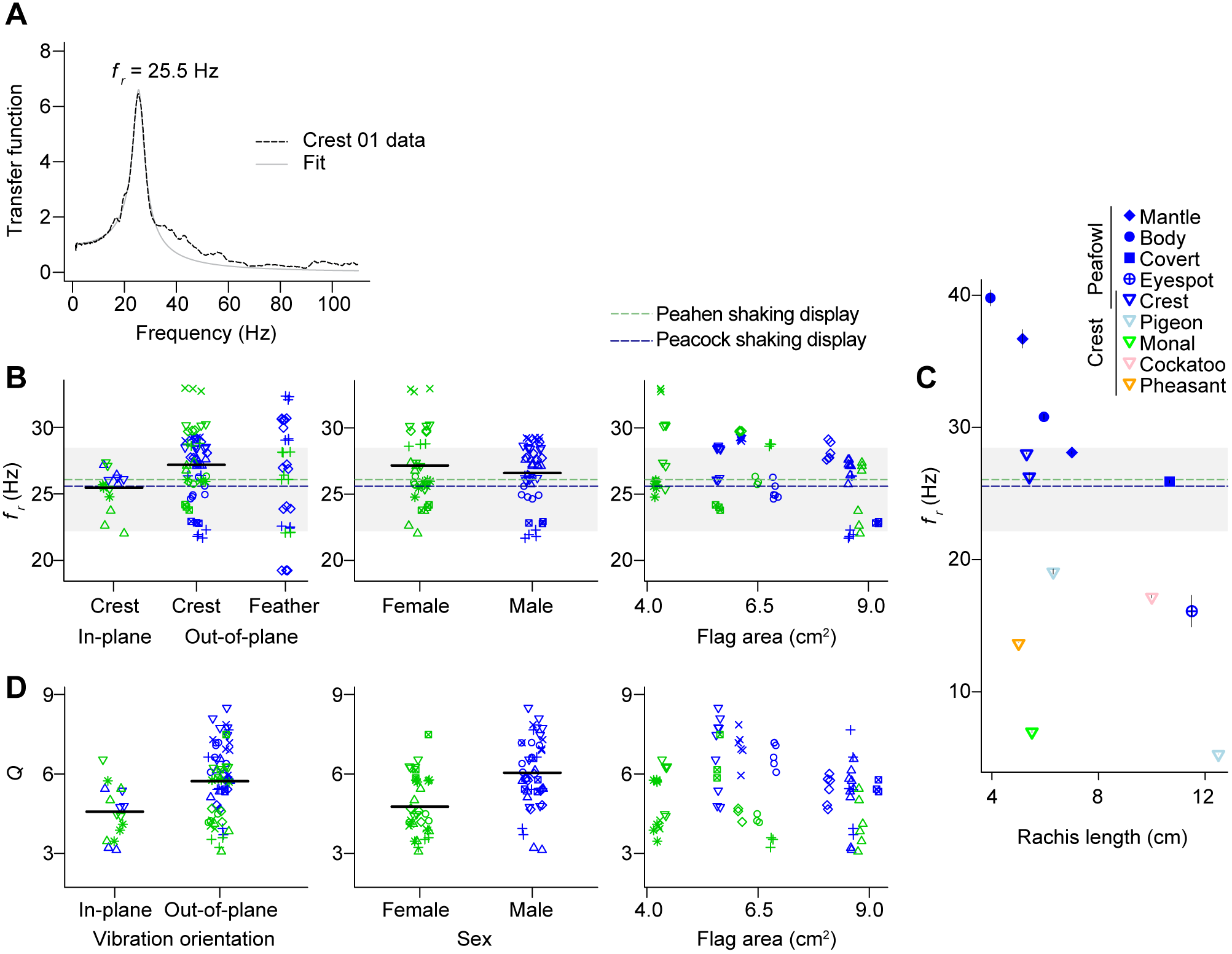
Vibrational resonance properties of peafowl crests and individual crest feathers. (A) Vibrational spectrum and Lorentzian fit for peacock crest sample Crest 01. (B-D) Data on the mean crest resonant frequencies, *f_r_*, and quality factors, *Q*. Each dried crest sample (n = 8 female, n = 7 male) is indicated by a unique symbol-color combination, consistent with Fig 2. (B) The mean resonant frequencies, *f_r_*, of the crest are a close match for the range of vibrational frequencies used during peafowl social displays. As an indication of measurement error, the average 95% CI for each mean *f_r_* estimate spans 0.072 Hz. The gray shaded area is the range of vibrational frequencies of the train-rattling display, with dotted lines showing the means for displays performed by peacocks (blue) and peahens (green) [20]. Variation in *f_r_* was influenced by the vibrational orientation and was also associated with the sex of the bird, but there was no significant association with the area of pennaceous flags at the top of the crest. The first panel in (B) also shows how a small sample of single crest feathers (n = 3 from male Crest 03, n = 5 from male Crest 05, and n = 3 from female Crest 10) had a similar range of resonant frequencies as the whole crests vibrated in the same out-of-plane orientation. (C) Fundamental frequency for vibrations in the out-of-plane orientation for peafowl crest and non-crest feathers with similar lengths and crest feathers from four non-peafowl species described in S1 Table and S1 Fig. Means for male and female peafowl crests are both plotted. The y-axis of (C) is aligned with that of (B) for comparison. (D) The mean quality factor, *Q*, was also influenced by the vibrational orientation, and was associated with the sex of the bird and the area of pennaceous flags. The average 95% CI for each mean *Q* estimate spanned 0.233. Black horizontal lines in (B) and (D) are grand means.

The repeatability of *f_r_* for whole crests was very high at 92% (95% confidence interval, 87-94%), demonstrating strong and consistent differences among individual crests (Fig 3B). Analysis of the sources of variation in *f_r_* indicated that 28% of the total variation in *f_r_* could be explained by sex, crest orientation, and the total area of the pennaceous flags (Fig 3B; see S5 Table for the best-fit model). The effect of crest orientation was strong and significant, such that out-of-plane vibrations have *f_r_* values approximately 2.4 Hz higher on average (p < 0.0001), whereas the sex difference was not significant (p = 0.86) and crests with reduced flag area have a slight but non-significant tendency to have higher *f_r_* values (p = 0.10). Crest length, width, number of feathers, and the percent of unaligned and short feathers did not explain variation among crests in the value of *f_r_*. The frequency response of individual crest feathers was generally consistent with that of the whole/intact crests, as the resonant frequencies of these feathers in the out-of-plane orientation ranged from 19.2 Hz to 32.4 Hz (Fig 3B).

Fig 3C compares the fundamental frequency of out-of-plane vibrations vs rachis length for peafowl crests and individual crest feathers, three other types of short peacock feathers, and crest feathers for four other species of birds. These data show that that the resonant frequencies of peafowl crests do not agree within measurement uncertainty with those of three other types of peafowl feathers, nor do they agree with those of crest feathers from four other bird species, even when the dependence of frequency on rachis length is taken into account.

Fig 3D shows that the mean quality factor *Q* for peafowl crests vibrated in the out-of-plane orientation (4.8 [4.0, 5.6] for females, 6.2 [5.6, 6.9] for males) was intermediate between that of peafowl eyespot feathers (*Q* = 3.6-4.5 ± 0.4 and 1.8 ± 0.3, for individual feathers and feather arrays, respectively) and the tail feathers that drive the shaking, for which *Q_1_* = 7.8 ± 0.5 [20]. This indicates that peafowl crests are moderately sharply-tuned resonators.

The repeatability of *Q* was estimated at 47% (95% confidence interval, 17-55%), indicating moderate differences among crests in *Q*. Approximately 49% of the variation in crest *Q* could be explained by sex, crest orientation, and the total area of the pennaceous flags (Fig 3D, see also S5 Table). Male crests were significantly more sharply-tuned than those of females (p < 0.005), and crests that had less flag area tended to be more sharply-tuned (p = 0.04). Peafowl crests also have more sharply-tuned resonance when they are vibrated out-of-plane (p < 0.0001) as compared to the in-plane orientation.

Note that the complete analysis of vibration data can be reproduced using data and code available at: https://doi.org/10.6084/m9.figshare.5451379.v5 [70].

### Force impulse experiments

When ring-shaped air vortices impacted the crests, the barbs responded with clearly visible motion on video with the average amplitude of motion at the flags of 9.4 [4.3, 14.4] mm (Fig 4A). Analysis of the crest vibrational motion vs time revealed an exponentially decaying sinusoidal response; the mean natural frequency, *f_o_*, measured by the force impulse method agreed to ≤ ± 0.4 Δ*f* of the value of *f_o_* predicted by Eq 2 using values of resonant frequency, *f_r_*, and *Q* measured using sinusoidal forces and frequency sweeps (Fig 4B). Thus, vortices cause the feather crest to vibrate at its natural frequency, with a decrease in amplitude of 13% after 0.2 s, the approximate period of peafowl wing-shaking displays.

**Fig 4.**
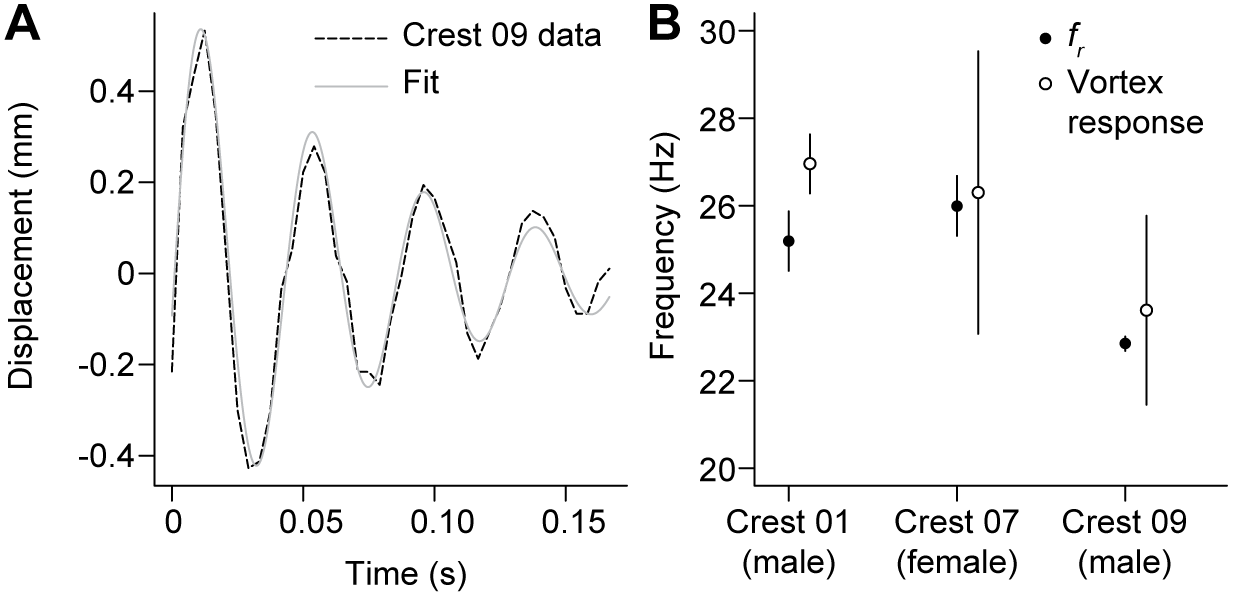
Displacement of the crest in response to air vortices. (A) Time series showing the change in flag position after a peacock crest (Crest 09) was impacted by a moving vortex of air. When peafowl crests were impacted by such air ring vortices, they deflected measurably, oscillating at their resonant frequency with an amplitude that decayed to a few percent of the initial value over the period of the peacock’s wing-shaking display. (B) Mean resonant frequencies (*f_r_*) and mean vortex response frequencies (± 95% CI) for three crests in the vortex experiment.

### Audio playback experiments

Fig 5A shows a waveform and spectrogram of a recording of train-rattling played back using the audio equipment in the playback experiment (see also S2 Fig). An example FFT power spectrum for the vibrational response of a peahen crest sample during audio playback is shown in Fig 5B. For train-rattling audio playback experiments in which the peahen crest samples were located in the flow near-field of the speaker, the vibrational power spectra of the samples had a peak well above noise near the playback train-rattling repetition rate (the effective drive frequency). However, when the white noise recording was played back, the spectral power near the drive frequency was < 4.3% of that found during playbacks. The peak frequency of crest vibrations agreed with the playback train-rattling repetition rate to within 95% CI for all measurements but one, for which it lay within 2.5 s.e.m. Measurements of crest vibrations made with an acoustic foam tile between the speaker and sample had < 11 % of the FFT spectral power at the drive frequency compared to measurements made without the foam; this value placed an upper bound on the contribution of background sources (e.g., room reverberations, substrate vibrations, etc.) that were not associated with particle-velocity oscillations from the playback stimulus.

**Fig 5.**
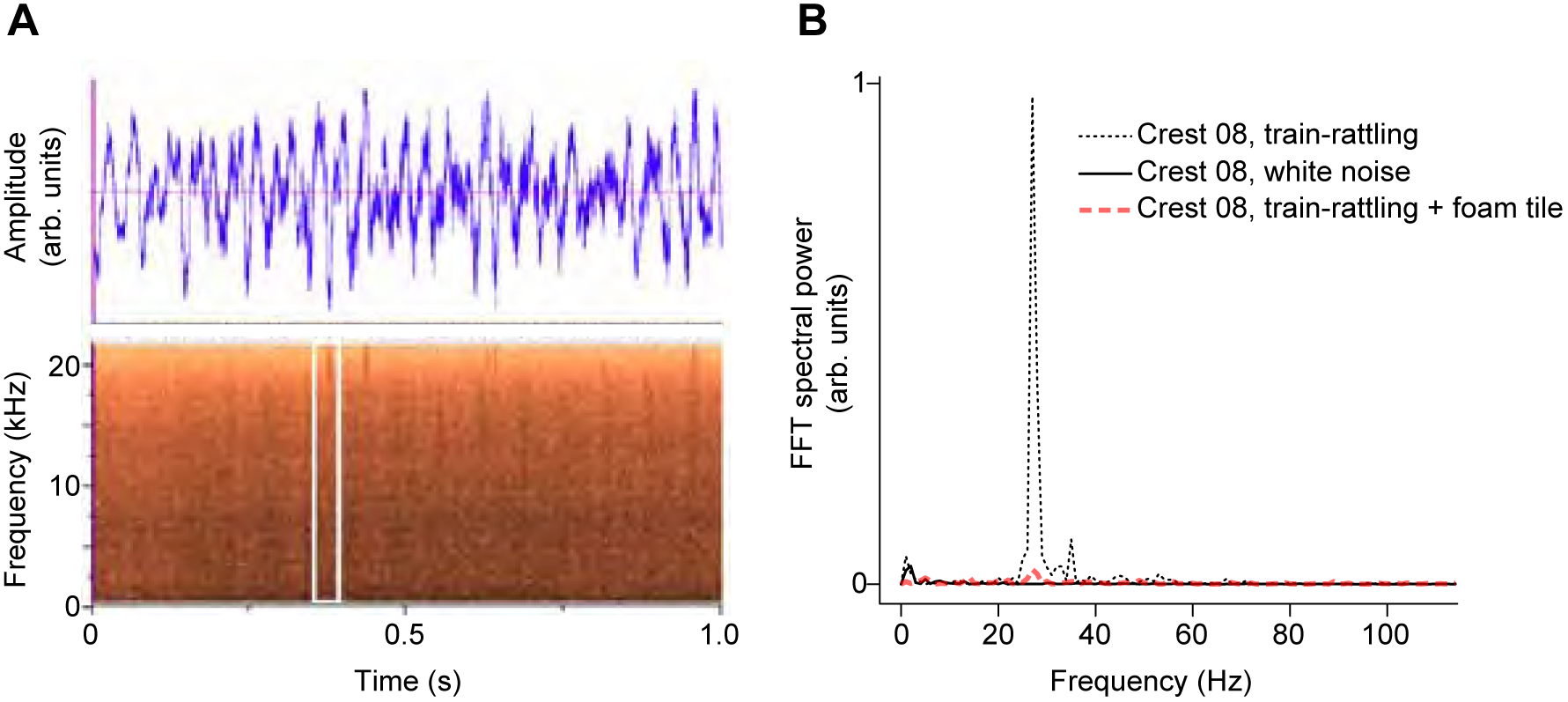
Effect of audio playback on crests. (A) An example waveform and spectrogram of the train-rattling sound used in the playback experiment. The white box in (B) highlights a single rattle note in the train-rattling spectrogram. (B) Vibrational response of a peahen crest (Crest 08) exposed to audio playback in the near-field of the speaker. The FFT spectral power during playback of train-rattling sound (dotted line, plotted on a linear scale on the y-axis) has a peak near the resonant frequency of the crest. The spectral power values recorded during white noise playback (solid line) and when the train-rattling audio was blocked by a foam tile (red dashed line) are also shown.

### Simulated wing-shaking experiments and wing-flapping during flight

The simulated wing-shaking experiment resulted in an airflow pattern with speeds ≤ 0.3 m/s. We used the measured positions of maximum airflow speed to determine the locations for three female crest samples for vibrational motion studies. The FFT power spectra of the crest flag vibrational motion had a single peak above the background noise at a frequency that agreed with the wing-shaking frequency within 95% CI (Fig 6) for distances up to 90 cm (one sample) and 80 cm (two samples) from the mean wingtip position.

**Fig 6.**
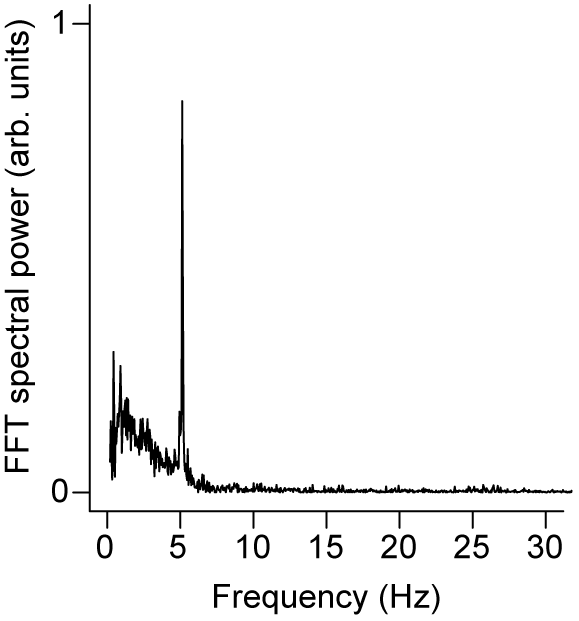
Effects of simulated wing-shaking displays. Vibrational response of a female peahen crest (Crest 13) exposed to airflow from a robot that simulated 5.0 Hz peacock wing-shaking displays at a distance 50 cm from the moving wingtip (see also S3 Fig). Note that the FFT spectral power (y-axis) is plotted on a linear scale.

The average peacock wing-flapping frequency during ascending and level flight was 5.5 [5.0, 6.1] Hz (S3 Table). This frequency agrees with the average frequency of 5.4 Hz (range of individual bird means = 4.5-6.8 Hz) found for wing-shaking display frequencies measured in the field [20].

### Mechanical bending properties

All feather crests exhibited a highly linear elastic response in the bending experiments: force and displacement were linearly related for displacements up to 10.1 [9.1, 11.0] mm (adjusted *R^2^* = 0.983 [0.978, 0.989]. This allowed us to compute the bending spring constant, k, from the fitted slopes (S4 Fig). The mean bending spring constants for the individual crests ranged from 0.0022 to 0.0054 N/mm with a measurement repeatability of 47% (95% confidence interval, 0-54%) due to the force sensor contacting the crest flag at somewhat different positions during different trials.

## Discussion

The fundamental vibrational resonant frequencies of peafowl crests were found to agree closely with the frequencies used during male train-rattling and female tail-rattling displays in Fig 3B. By contrast, these display frequencies do not agree with the resonant frequencies found for feathers of similar length from other parts of the peafowl’s body, or with those found for the crest feathers of four other bird species in Fig 3C, which collectively span a frequency range that is nearly seven times that of the observed range of rattling displays. This means that the close frequency match between peafowl displays and crest resonance is not due simply to species, type of feather (i.e., crest vs. tail), or rachis length. This finding agrees with prediction P2 (Box 1) that crest feathers with a mechanosensory function would have a frequency response tuned to match stimuli with a well-defined frequency.

Our results also indicate that both the resonant frequency and the *Q* factor of the peafowl crest’s vibrational response should agree with those of the array of tail and train feathers that produce the shaking display, which have been previously characterized in [20]. This implies that the peafowl’s crest would be well-matched to the train’s mechanical sound, but not to environmental sources of noise [31]. In agreement with prediction P4 (Box 1), we also found that exposing peahen crest samples to the near-field of audio playbacks of train-rattling sounds caused the crests to vibrate detectably on video at their resonant frequency (Fig 5B). By contrast, exposing crest samples to white noise resulted in no measurable vibrations above background noise levels. We therefore hypothesize that this match of vibrational resonant properties might have functional significance during multimodal courtship displays that generate mechanical sound.

As found for live birds [17], the peafowl crest samples had relatively uniform lengths and numbers of feathers (Fig 2). While our crest samples had slightly lower flag area than fully spread crests of living birds, we found that individual crest feathers had similar vibrational responses to those of entire crests, indicating that interactions between crest feathers is not the main determinant of resonant frequency. This indicates that the results of our vibrational dynamics experiments are also applicable to crests *in vivo*, on the live bird.

Peafowl crests do not have a resonant frequency near the 5.4 Hz rate of wing-shaking displays. However, we still found that peafowl crests vibrated detectably in response to impulsive airflows similar to those produced during wing-shaking (prediction P3; Box 1). By measuring the deflection of peafowl crests when they were struck by individual air ring vortices (Fig 4), we found that each impulsive stimulus generated a distinct crest response in which the crest feathers briefly vibrated at their natural frequency, before decaying to zero in a time comparable to that of the wing-shaking period. This result provided independent validation of the resonant frequency of crests, measured from the spectral responses in Fig 3. It also means that periodic but isolated force impulses generated by the wing-shaking display are effectively experienced as distinct stimuli that cause the crest to oscillate only briefly near resonance, akin to an infrequently struck bell. Further confirming this interpretation, we found that airflows due to simulated wing-shaking at distances from the crest ≤ 90 cm drove measurable transient crest deflections (Fig 6), similar to the minimum male-female distances in the field during such displays. The linearity of the measured elastic response also suggests that this result can be extrapolated to greater distances. These findings imply that the airflow impulses generated by *in vivo* wing-shaking displays could stimulate the feather crests of nearby female by producing a series of distinct vibrational responses.

Our measurements of vibrational responses during audio playback were limited to relatively large amplitudes and small distances compared to the very small thresholds found for other mechanoreceptors *in vivo*, and consequently to relatively small source-receiver distances. However, the low thresholds found for mechanosensation *in vivo* suggest that the actual detection range could be much greater than our *in vitro* limits. For example, pigeons can detect submicron threshold vibrational amplitudes applied to flight feathers [93,94], mammalian hair cells are sensitive to sub-nanometer displacements and 0.01 deg rotations [95], tactile receptors in human skin are sensitive to submicron vibrational amplitudes [96], and insect filiform hairs are sensitive to airspeeds as low as 0.03 mm s^−1^ [97]. This idea also is supported by our measured linear elastic response of feather crests to bending (S4 Fig), which indicates that our results can be extrapolated linearly to lower magnitude stimuli corresponding to larger source-sample separation than those measured here during the audio playback experiments. Our microscopy results confirmed that peafowl crest feathers have feathers at their bases with the morphology expected for filoplumes (prediction P1; Box 1). However, further histological and electrophysiological studies of the receptors at the base of avian crest feathers and their associated filoplumes are needed to determine whether these crests can in fact function as sensors that are sensitive to airborne stimuli like the ones studied here.

Given that feathers are known to function as airflow sensors during flight, it is easy to imagine how they also could be adapted to function as sensors during social signaling. For example, during social displays, many birds flap or vibrate their wings or tails [66,67,98,99,20], producing dynamic visual stimuli as well as mechanical sound and periodic air flow stimuli. Thus, these multimodal displays have the potential to stimulate multiple senses, including vision, hearing, and vibrotactile perception. Several non-exclusive scenarios could provide a functional benefit of such a close frequency match. For example, we can hypothesize that air-borne stimuli generated by train-or tail vibrating or shaking displays provide females with an indication of male muscle power and endurance, or that stimuli generated by wing-shaking displays serve as signals of flight muscle performance [99]. Another hypothesis is that male displays have been selected to match and stimulate pre-existing mechanosensory properties of the female crest, without any benefits to females of this close match. Yet another hypothesis is that both males and females experience crest vibrations driven by body oscillations due to their own train-and tail-rattling displays as a form of proprioception. Conversely, our measurements do not support a visual display function for peafowl feather crest vibrations because the resulting amplitudes of a few mm at most are unresolvable given limitations due to the peafowl’s visual acuity [20].

Although we have demonstrated here that peafowl crest feathers are effectively stimulated by airborne stimuli during social displays, we do not yet know whether this has behavioral or social significance. Testing this hypothesis requires *in vivo* behavioral experiments. Crest vibrations are challenging to measure directly, given that both sexes move frequently during displays and are viewed against complex visual backgrounds. Instead, a first step in peafowl could be to blindfold females and test whether airborne stimuli at the socially salient frequencies elicit a behavioral response. Further experiments could test the function of the crest during male courtship displays by removing or altering the female crest and then examining how females respond to male displays. One way to do this would be to apply a thin coat of clear varnish to the rachis of the crest feathers; this would stiffen the rachis and increase resonant frequency without affecting the crest’s visual appearance (i.e., size or flag iridescence). Similar manipulations could also test whether peacocks use proprioception from the crest to modulate their own vibration displays. The movement of females during displays could also be examined in relation to the airflow patterns generated by wing-shaking peacocks, to test whether female movements are correlated with specific airflows generated by the males. These correlative results could then be tested experimentally by measuring the behavioral response of peahens to oscillatory air flows modulated at frequencies close to and distinct from their crest resonance, to see how this influences attention and body orientation. Because audible sound cues are omnidirectional, these responses could be distinguished from hearing-induced behaviors by comparing results when air flows are directed toward and away from specific regions of the birds’ plumage.

The biomechanical properties of the peafowl’s crest also suggest a novel design for making sensitive biomimetic detectors for sensing impulsive or periodic airflows. Such devices are required for proposed robotic applications of air vortex rings and other airflow signals as a communication channel [100]. The addition of an extremely lightweight pennaceous flag to a cantilever made from a resistance-based flex sensor would enable the flex sensor to experience a large torque from a small force, with a minimal increase in mass.

Thus far, the elaborate shape, size and color of many bird feather crests has led to an emphasis on their visual appearance [14]. However, many avian courtship displays also involve wing-shaking, tail-fanning and mechanical sound production that may be detected by nearby females in the vibrotactile channel. For example, we have compiled a list of at least 35 species distributed in 10 avian orders that have crests and perform these types of displays (S6 Table). Given the growing interest in multisensory signaling, it seems worth pursuing behavioral studies to investigate whether mechanosensory stimulation enhances the reception of visual and acoustic cues during this multimodal display. The close match between the resonant frequencies found here for peafowl crests and this species’ social displays suggest that it is time to explore the hypothesis that birds receive and respond to vibrotactile cues in a wider variety of scenarios.

## Acknowledgments

We are grateful to Maarten Hesseling for assistance with preliminary vibrational measurements, Robert Beyer, Robert Lukasik, and Roger Hill for help with instrumentation design and construction, Kate Davidson (Siskiyou Aviary) and James Hare for feather samples, and Robert Koch, Holger Klinck, and Carr Everbach for advice about reproducing low repetition rate sounds. We thank two anonymous reviewers and James Hare for helpful comments on an earlier version of the manuscript.

## Author contributions

Conceptualization: S.A.K.; Methodology: S.A.K., D.V.; Investigation: S.A.K., D.V.; Data curation: S.A.K., D.V., R.D.; Analysis: S.A.K., R.D., D.V.; Writing – original draft: S.A.K., R.D.; Writing – review & editing: S.A.K., R.D., D.V.

## Data availability

All data and code necessary to reproduce the results of this study are available at: https://doi.org/10.6084/m9.figshare.5451379.v5

---

**S1 Fig.**
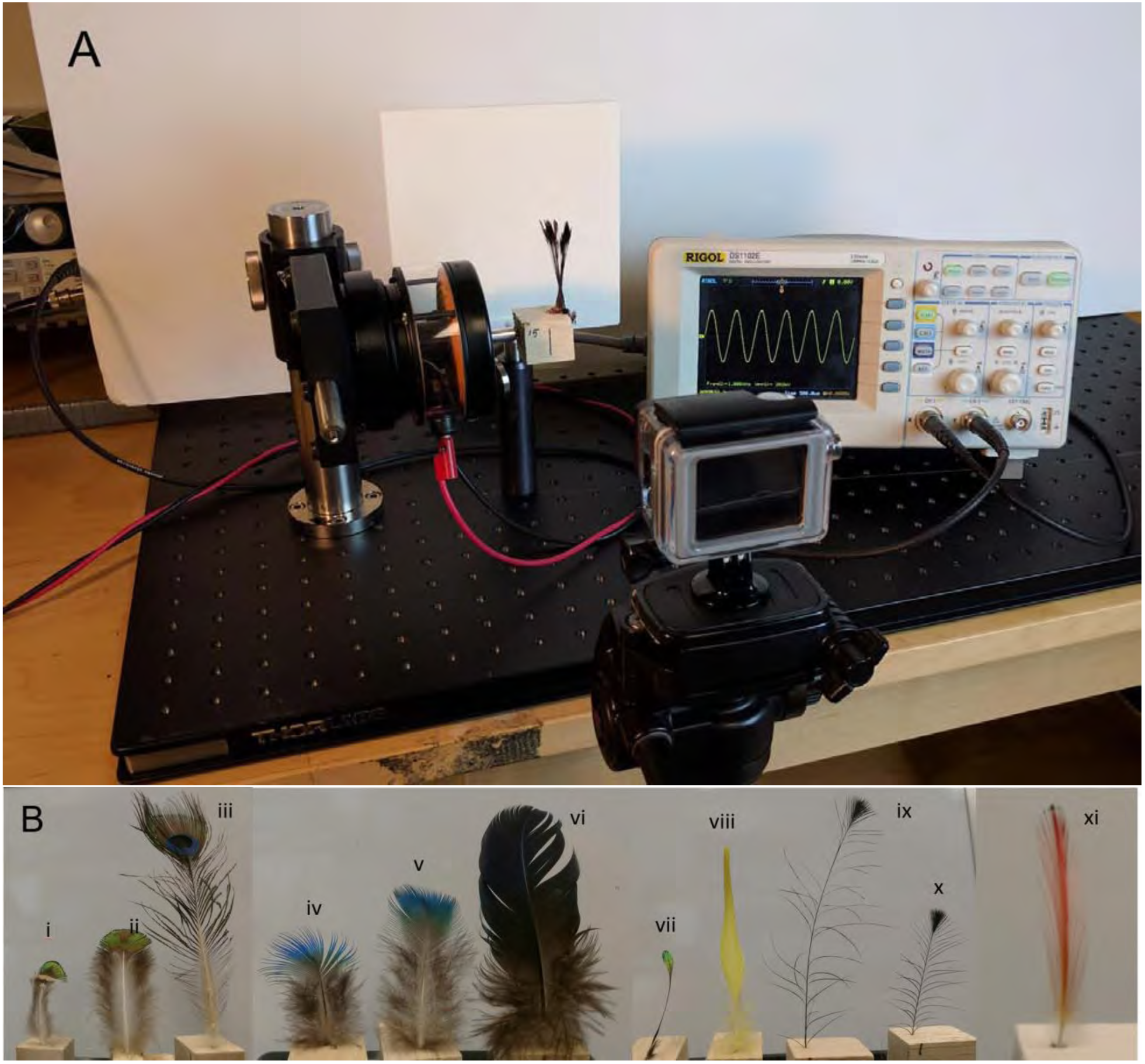
Vibrational response apparatus and feather samples. (A) Apparatus for measuring the vibrational response of peafowl feather crests. Crests were first glued onto balsa wood blocks, which were then mounted on a mechanical shaker driven by a function generator that produced a sine wave output with a linear ramp in the frequency of shaking. The resulting motions of the crest flags and the shaker were measured using high-speed video. (B) Feather samples from Table S1 measured for comparison with peafowl crest vibrational resonant frequencies (not shown to scale): (i), (ii) peacock mantle feathers; (iii) short peacock eyespot feather; (iv), (v) peafowl body semiplumes; (vi) peacock wing covert; (vii) Himalayan monal crest feather); (viii) yellow crested cockatoo crest feather; (ix), (x) Victoria crowned pigeon crest feathers; (xi) golden pheasant crest feather.

**S2 Fig.**
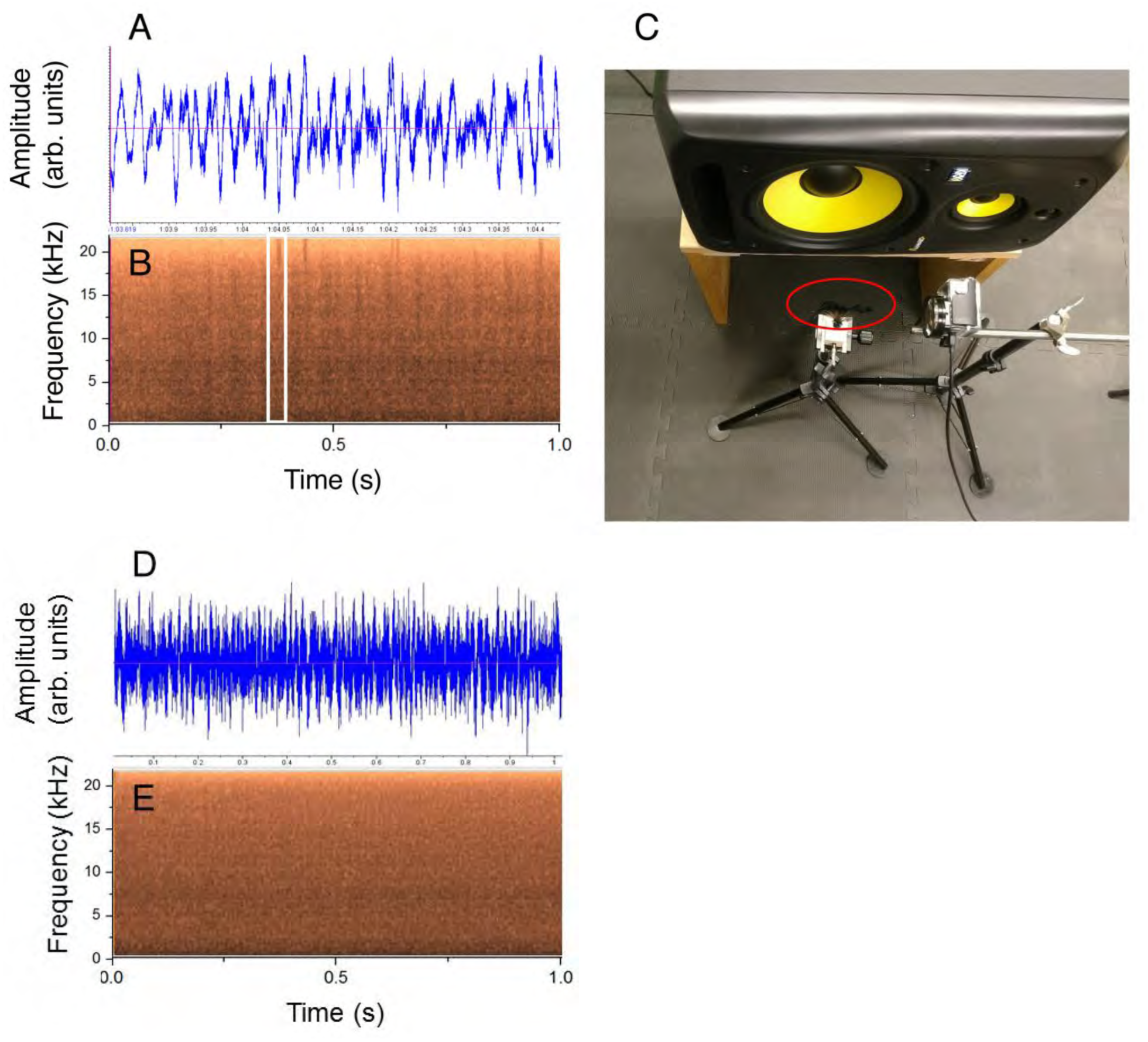
Example audio and apparatus used for measuring the vibrational response of peafowl feather crests during audio playback of peacock train-rattling mechanical sounds. (A-B) An example waveform and spectrogram for one of the playback stimulus tracks of peacock train-rattling sounds. The waveform (A) and spectrogram (B) were generated from a re-recording made of the playback stimulus, to ensure that features of the playback stimulus matched those of the original recording from Dakin et al. (2016). The white box in (B) highlights a single rattle note in the train-rattling spectrogram. (C) Playback apparatus viewed from above. The crest sample (red ellipse) was exposed to the flow near-field of a loudspeaker (top) that played back peacock train-rattling sounds. The resulting motions of the crest flags were measured using a high-speed video camera (located to the right of the ellipse). (D-E) Waveform and spectrogram of the white noise control played back through the same audio system used for train-rattling playbacks. This illustrates the resemblance between the broad-band frequency spectrum of the white noise control (E) and the rattle notes (B). However, the white noise control lacks modulation at the low frequencies characteristic of displays.

**S3 Fig.**
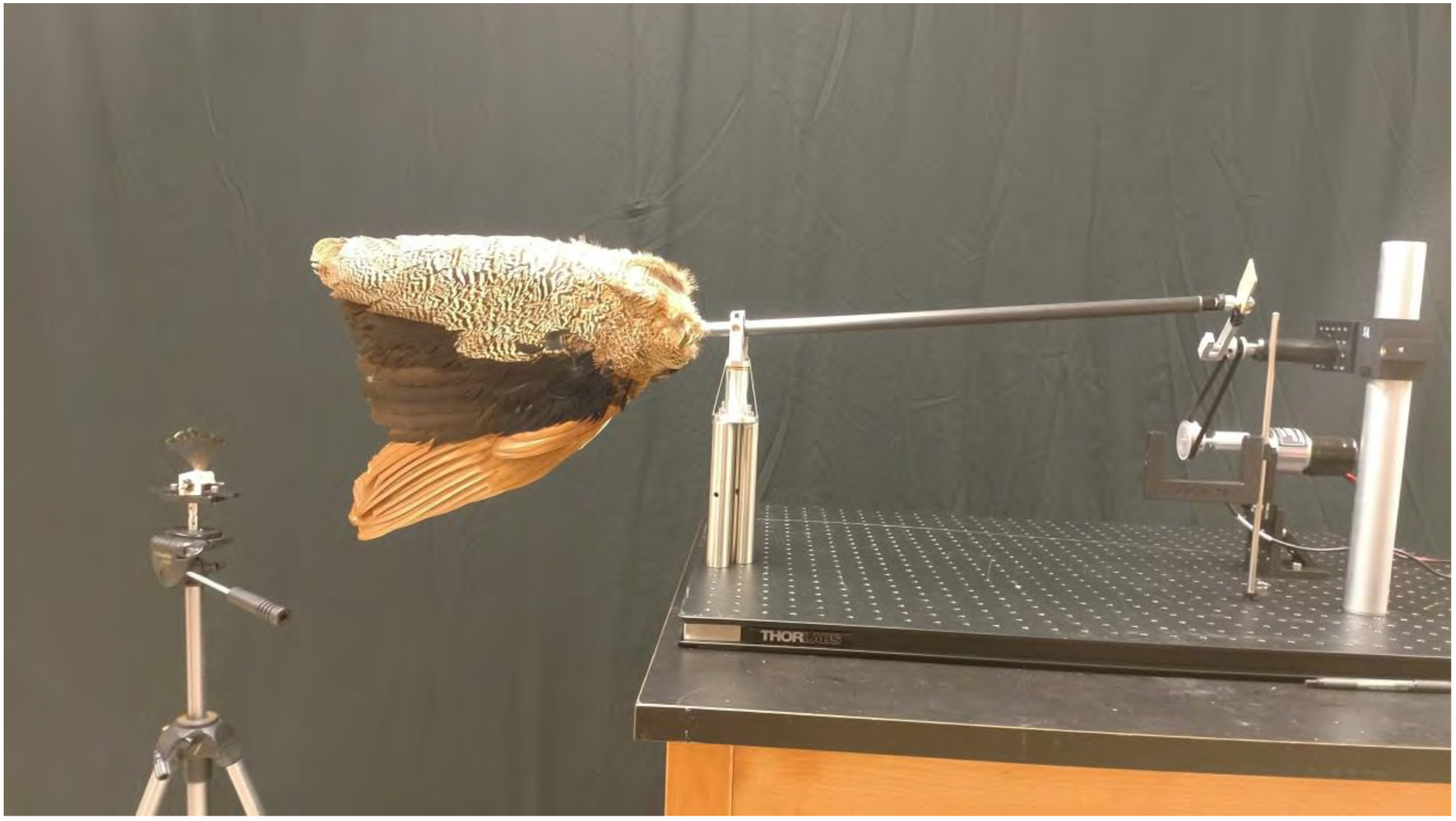
Robotic apparatus for simulating peacock wing-shaking. Peacock wing-shaking displays were simulated using a peacock wing mounted on a carbon fiber rod. The rod was rotated at approximately 5 Hz (a typical wing-shaking frequency) about a clevis joint located at the wing’s shoulder joint, ensuring that the plane of the wing’s surface remained vertical while the tips circumscribed a 14 cm diameter circle. Peahen crests were positioned in the region of maximum airflow at distances ≤ 90 cm (50 cm shown here) from the wingtips. The resulting crest motion was filmed using high-speed video.

**S4 Fig.**
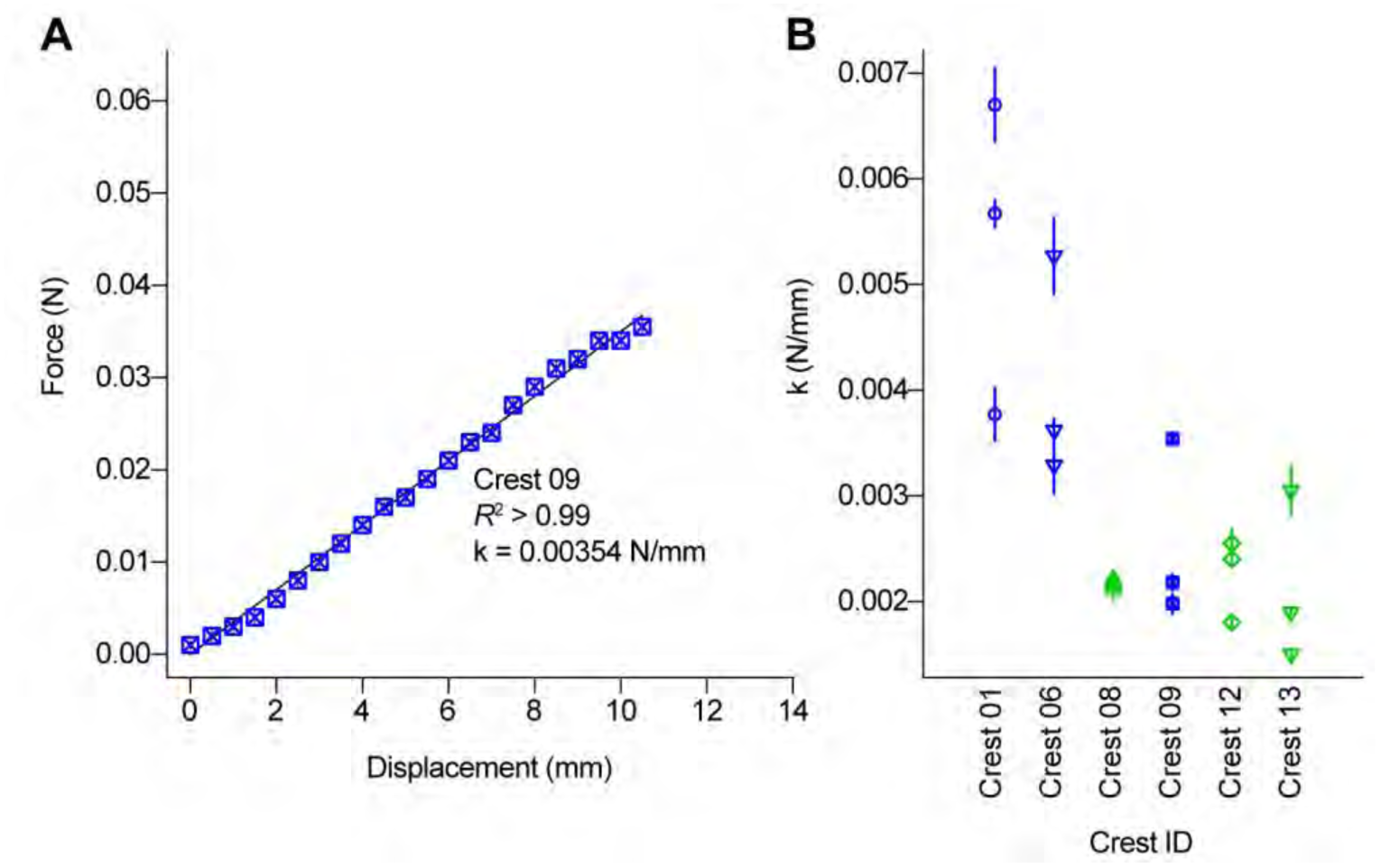
Bending spring constant, k, of peafowl crests. Force-displacement trials were performed three times each for n = 3 male and n = 3 female crests, respectively. The bending spring constant, k, was calculated from the slope of linear model fits to the resulting force-displacement data from each trial. The example in (A) shows data from a single trial on peacock Crest 09 to illustrate the linearity of the response, with symbols scaled to span y-axis measurement error. (B) Values of k from each of three trials on the total n = 6 crests. Each crest sample is denoted by a different symbol-color combination, following Figs. 2-3 of the main text, and ordered left to right by decreasing mean k value. Blue data are male (peacock) crests and green data are female (peahen) crests.

**S1 Table.** Individual feathers used in the vibrational resonant measurements. These values are provided for comparison with data shown in Fig. 3 for peafowl crests.

| Species | Feather type | Rachis length (cm)<br>(+/- 0.05 cm 95% CI) |
| --- | --- | --- |
| Indian peafowl male | Body contour semiplume | 3.94 |
| Indian peafowl male | Body contour semiplume | 5.95 |
| Indian peafowl male | Wing covert | 10.67 |
| Indian peafowl male | mantle | 5.15 |
| Indian peafowl male | mantle | 7.00 |
| Indian peafowl male | train eyespot | 11.50 |
| Victoria crowned pigeon | crest | 6.30 |
| Victoria crowned pigeon | crest | 12.50 |
| Himalayan monal | crest | 5.50 |
| Golden pheasant | crest | 5.00 |
| Yellow-crested cockatoo | crest | 10.00 |

**S2 Table.** Sound pressure levels (SPL) of avian wingbeats during flight and wing-beating displays measured in previous studies.

| Bird species | SPL (dB) | Distance (m) | Reference |
| --- | --- | --- | --- |
| Eastern phoebes,<br>*Sayornis phoebe* | 64-66 dB SPL at 1 kHz | 1.2 m | Fournier, J.P., Dawson, J.W., Mikhail, A., & Yack, J.E. (2013). If a bird flies in the forest, does an insect hear it? *Biology letters* 9(5): 20130319. |
| Black-capped chickadees,<br>*Poecile atricapillus* | 54-60 dB SPL at 25 kHz | 1.2 m | Fournier, J.P., Dawson, J.W., Mikhail, A., & Yack, J.E. (2013). If a bird flies in the forest, does an insect hear it? *Biology letters* 9(5): 20130319. |
| Crested pigeons,<br>*Ocyphaps lophotes* | $\leq 67.6$ dB SPL | 1.0 m | Hingee, M. & Magrath, R.D. (2009) Flights of fear: a mechanical wing whistle sounds the alarm in a flocking bird. *Proceedings of the Royal Society of London B*: rspb20091110. |
| Ruffed grouse,<br>*Bonasa umbellus*<br>(a wing-beating display) | 66.2 dB SPL, bandwidth 300 Hz to 8 kHz, frequency weighting not reported | 1.0 m | Garcia, M., Charrier, I., & Iwaniuk, A.N. (2012) Directionality of the drumming display of the ruffed grouse. *The Condor* 114(3): 500-506. |

**S3 Table.** Wingflap frequencies of adult peacocks during level and ascending flight.

| Source | Number of individuals | Wingflap frequency (Hz) |
| --- | --- | --- |
| https://www.youtube.com/watch?v=HvY_1wFSFsQ<br>accessed September 28, 2017 | 1 | 4.92 |
| https://www.youtube.com/watch?v=7gxQwm4MWns<br>accessed September 28, 2017 | 1 | 4.16 |
| https://www.youtube.com/watch?v=U55iMIiI_k0<br>accessed September 28, 2017 | 1 | 6.43 |
| https://www.youtube.com/watch?v=kZe0jLkeMuk<br>accessed September 28, 2017 | 1 | 4.42 |
| https://www.youtube.com/watch?v=FrMQs7OwWC8<br>accessed September 28, 2017 | 3 | 6.36<br>6.00<br>5.55 |
| https://www.youtube.com/watch?v=A5xSgaXDkTY<br>accessed September 28, 2017 | 2 | 5.81<br>6.15 |

**S4 Table.** Symbols used to indicate different crest samples in Figures 2 and 3 of the main text.

| Symbol | Sample ID | Sex |
| --- | --- | --- |
| ○ | Crest 07 | female (peahen) |
| △ | Crest 08 | female |
| + | Crest 10 | female |
| × | Crest 11 | female |
| ◇ | Crest 12 | female |
| ▽ | Crest 13 | female |
| ■ | Crest 14 | female |
| * | Crest 15 | female |
| ○ | Crest 01 | male (peacock) |
| △ | Crest 02 | male |
| + | Crest 03 | male |
| × | Crest 04 | male |
| ◇ | Crest 05 | male |
| ▽ | Crest 06 | male |
| ■ | Crest 09 | male |

**S5 Table.** Best-fit models of *f_r_* and *Q* in the analysis of the vibrational dynamics measurements.

| Response | Fixed-effect | Estimate (SE) | t | p |
| --- | --- | --- | --- | --- |
| $f_r$ | Orientation (in-plane vs. out-of-plane) | −2.22 (0.23) | −9.46 | < 0.0001 |
|  | Sex (male vs. female) | 0.26 (1.49) | 0.18 | 0.86 |
|  | Top area | −0.82 (0.46) | −1.78 | 0.10 |
| $Q$ | Orientation (in-plane vs. out-of-plane) | −1.26 (0.25) | −4.94 | < 0.0001 |
|  | Sex (male vs. female) | 1.85 (0.49) | 3.71 | 0.005 |
|  | Top area | −0.37 (0.15) | −2.40 | 0.04 |

**S6 Table.**
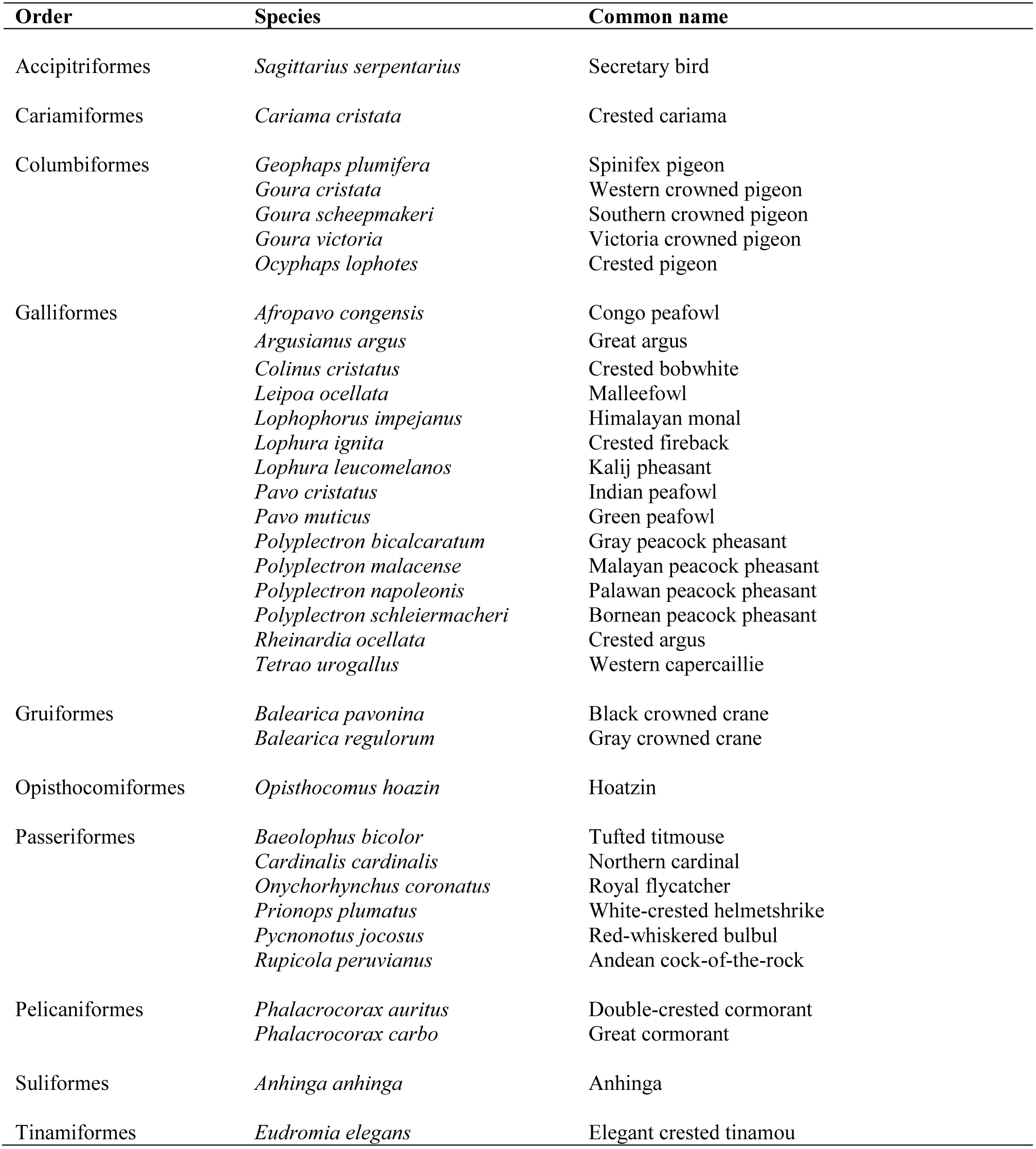
Species in which both sexes have crests of flexible feathers and the male also performs a shaking display. There are many bird species wherein both sexes have a flexible feather crest. To understand the taxonomic breadth of birds that have shaking displays in addition to the crest, we used natural history resources including photos, videos and descriptive accounts of appearance and behavior. We documented at least 35 species across 10 different orders in which the females exhibit flexible feather crests and the males are known to perform shaking displays.

